# The MS remyelinating drug bexarotene (an RXR agonist) promotes induction of human Tregs and suppresses Th17 differentiation *in vitro*

**DOI:** 10.1101/2021.02.17.431344

**Authors:** Christof Gaunt, Daniel Rainbow, Ruairi Mackenzie, Lorna Jarvis, Hani Mousa, Nicholas Cunniffe, Zoya Georgieva, J. William Brown, Alasdair Coles, Joanne Jones

## Abstract

The retinoid X receptor (RXR) agonist bexarotene has recently been shown to promote remyelination in individuals with multiple sclerosis. Murine studies demonstrated that RXR agonists can have anti-inflammatory effects by enhancing the ability of all-trans-retinoic acid (*αt*RA), the primary active metabolite of vitamin A, to promote T regulatory cell (Treg) induction and reduce Th17 differentiation *in vitro*, following stimulation of naïve CD4 cells in the presence of TGF-*β*.

Stimulating naïve human CD4 T cells for 7 days, in the presence of either Treg or Th17 skewing cytokines ± bexarotene (1 μM), ± other RXR agonists (9*Cis*RA and NRX 194204), or ± *αt*RA (100 nM) shows that RXR agonists, including bexarotene, are capable of tipping the human Treg/Th17 axis in favour of Treg induction. Furthermore, this occurs independently of *αt*RA and retinoic acid receptor (RAR) signalling. Tregs induced in the presence of bexarotene express many of the canonical markers of T cell regulation and are functionally suppressive *in vitro.*

These findings support a potential immunomodulatory role for bexarotene and highlight the possible therapeutic application of RXR agonists in autoimmune disease, with bexarotene’s pro-remyelinating effects making multiple sclerosis a particularly attractive disease target.

**Significance Statement:** The pan-retinoid X receptor (RXR) agonist bexarotene has recently been shown to promote remyelination in patients with multiple sclerosis. Here we demonstrate that bexarotene, and other RXR agonists have immunomodulating effects, tipping the Th17/T regulatory cell (Treg) differentiation axis in favour of Treg development.

These findings lend support to the idea of developing RXR agonists as treatments of autoimmune diseases, in particular multiple sclerosis.

## Introduction

Bexarotene, a pan retinoid X receptor (RXR) agonist licensed for use in humans as a treatment of cutaneous T-cell lymphoma, has recently been shown to promote remyelination in multiple sclerosis (MS). Although the Cambridge Centre for Myelin Repair (CCMR)-one trial failed to meet its primary endpoint (reported at the 2020 ACTRIMS/ECTRIMS conference), pre-planned analyses demonstrated that bexarotene promoted the remyelination of the most demyelinated lesions, particularly in grey matter (as determined radiologically). In addition, conduction in the visual pathway improved in patients randomised to bexarotene, providing further evidence of its pro-remyelinating effect.

The CCMR-One clinical trial largely came about because of work done in rodents, which demonstrated that oligodendrocytes express RXR (specifically RXR-⋁) during active remyelination, and that remyelination can be promoted, *in vitro* and *in vivo*, following administration of RXR agonists [2].

Murine studies have also demonstrated that RXR agonists can enhance the ability of all-trans-retinoic acid (*αt*RA), the primary active metabolite of vitamin A, to promote Treg induction (iTregs), and reduce Th17 differentiation *in vitro*, following stimulation of naïve CD4 cells in the presence of TGF-*β* [3]. At physiological concentrations, *αt*RA binds to the retinoic acid receptor (RAR) which is bound to DNA as a heterodimer with RXR in regions called retinoic acid response elements (RAREs). Binding of *αt*RA to the RAR/RXR heterodimer leads to a conformational change, which in turn promotes down-stream transcription of hundreds of genes [4], including those involved in the induction of Tregs.

In contrast to *αt*RA, RXR ligands are unable to promote the differentiation of murine naïve T cells to iTregs directly. Instead, they act synergistically with *αt*RA to promote FOXP3 expression [3]. In this way RXR is said to act as a conditionally permissive binding partner to RAR – that is, whilst RXR ligand binding is unable to trigger down-stream signalling, maximal receptor signalling and transcriptional activity is achieved when RAR and RXR ligands bind simultaneously [3, 4]

Here we sought to determine if the RXR agonist bexarotene, licensed for use in humans and recently shown to have pro-remyelinating effects in MS, has similar immunoregulatory effects on the human iTreg/Th17 axis.

## Results

### Bexarotene promotes the induction of human Tregs independent of RAR activation

To determine the effect of bexarotene on human iTreg induction FACS-sorted naïve CD4 T cells were cultured for 7 days in serum-free, and therefore *αt*RA free media, under iTreg inducing conditions (anti-CD3/28 stimulation plus IL-2 and TGF-β) with and without the addition of bexarotene (1 *μ*M), *αt*RA (100nM) or both. As a negative control, naïve cells were also stimulated with anti-CD3/28 beads in the presence of IL-2 only (“IL-2 control”).

In contrast to what was expected from murine studies, bexarotene alone was sufficient to increase Treg induction as measured by FOXP3 expression (Figure 1A, from 21.52% to 39.25% p=0.0038). This increase was equivalent to that seen following the addition of *αt*RA (41.29% p=0.0013). No synergistic effect between bexarotene and *αt*RA was observed (Figure 1B).

**Figure 1.**
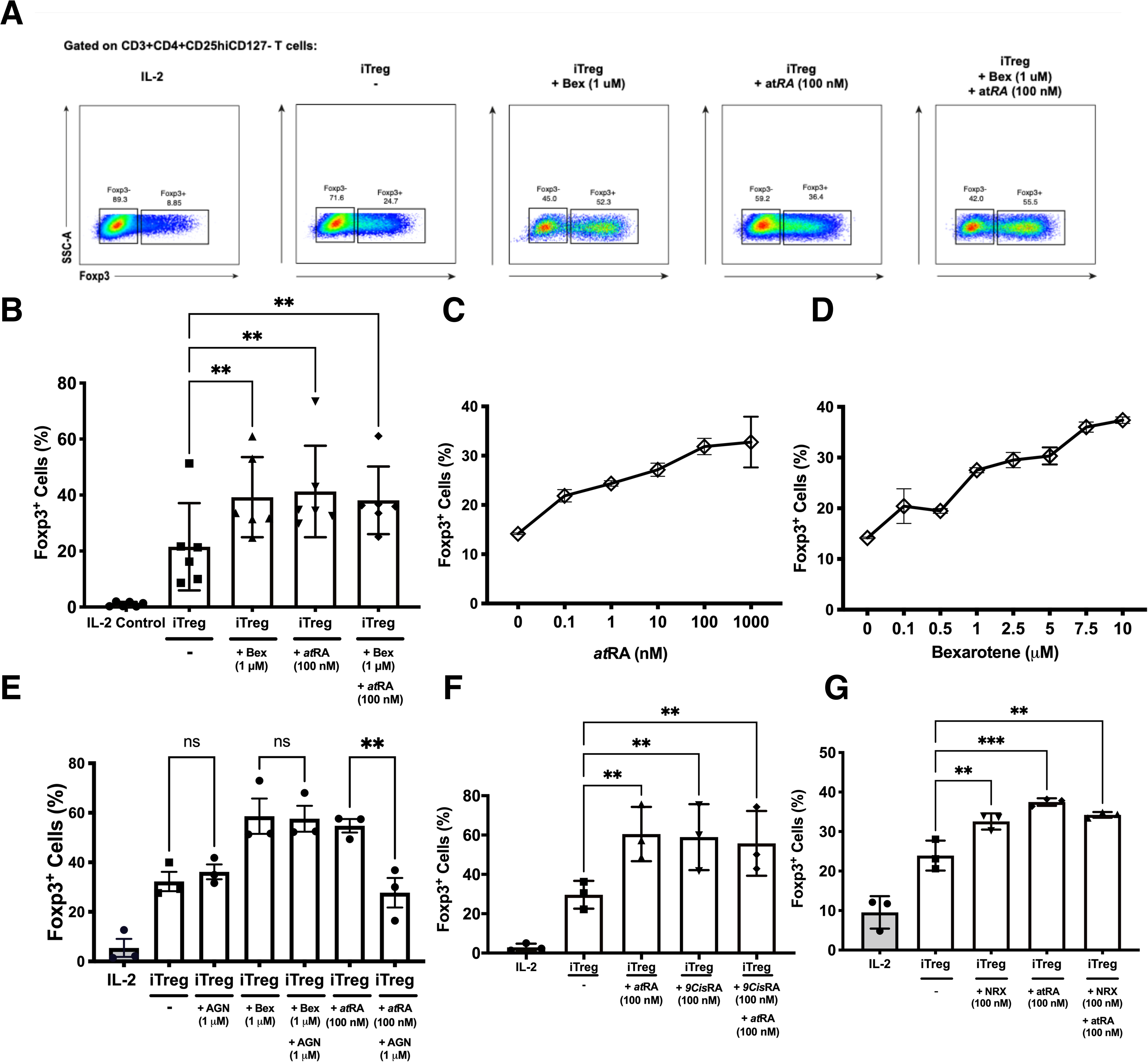
RXR agonists promote human iTreg differentiation independent of RAR signalling. (A) Gating strategy and example dot plots showing iTreg induction following culture of naïve T cells for 7 days in iTreg skewing conditions +/− bexarotene (1 μM), +/− atRA (100 nM) or both (B) Summary data for n=6 experiments (C-D) percent iTreg induction over a range of atRA and bexarotene concentrations. (E) Co-culture with the RAR antagonist AGN reverses increased iTreg induction seen in the presence of atRA, but not Bexarotene. (F-G) The RXR agonists 9CisRA and NRX194204 also promote human iTreg differentiation in vitro. Shown are mean values ± SEM. Significance calculated using a repeated measures one-way ANOVA. Šidák’s correction was performed.*p<0.05, **p<0.01, ***p<0.001 ns: not significant.

Next a dose ranging experiment was performed which demonstrated that bexarotene and *αt*RA caused a similar dose-dependent increase in iTreg induction, resulting in up to ~2.5 fold more FOXP3+ cells (Figure 1C,D)). At concentrations above 20 *u*M bexarotene inhibited iTreg differentiation and resulted in significant cell death (>95% dead; data not shown).

Although all assays were performed in serum-free media, and therefore *αt*RA free conditions; to be certain that bexarotene was not driving Treg differentiation via a RAR dependent mechanism, the above assays were repeated following pre-incubation with AGN 193109, a potent and selective RAR antagonist (Figure 1E). Pre-incubation with AGN fully blocked the pro-iTreg inducing effects of *αt*RA (% iTregs 54.78% vs 22.75%, p=0.0066, compared to iTreg conditions alone = 32.27%). In keeping with its RAR-independent mode of action, the addition of AGN had no effect on increased iTreg differentiation in the presence of bexarotene (% iTregs 58.63% vs 57.60%). AGN did not affect cell viability at the concentrations used (data not shown).

### 9-*cis*RA and the synthetic RXR agonist, NRX 194204 also promote human iTreg differentiation *in vitro*

To determine if the effects seen were specific to bexarotene, or a generic effect of RXR agonism, FACS sorted naïve T-cells were cultured in serum-free media under iTreg inducing conditions (IL-2 and TGF-β) with and without the addition of a synthetic RXR agonist (NRX 194294) or 9-*cis*RA (a metabolite of Vitamin A, and the primary endogenous RXR agonist).

As seen with bexarotene, both 9-*cis*RA and NRX enhanced to the same degree naïve T-cell differentiation to iTregs independently of RAR agonism, and neither had a synergistic effect with *αt*RA (% iTregs with 9-cisRA= 58.93, % iTregs with NRX= 32.56%; Figure 1,F,G).

### Bexarotene induced Tregs are suppressive *in vitro*

In addition to expressing FOXP3, the regulatory phenotype of the iTregs was confirmed through suppression assays. In brief, naïve CD4+ T cells were isolated from healthy donors and following 7 days of iTreg induction protocol with and without bexarotene, cells were co-cultured with Teffectors for 3 days and proliferation measured.

Tregs induced in the presence of bexarotene were as suppressive as Tregs induced in standard Treg induction conditions (59.2% suppression at a ratio of 1:6, Teff:iTregs) (Figure 2A). This suppressive effect was demonstrated to be titratable. At ratios of 1:1, 3:1 and 6:1, there was a non-statistically significant trend for iTregs induced in the presence of bexarotene to be more suppressive than iTregs induced without bexarotene (+9.7%, +13.1% and +7.1% respectively). However, this is likely to reflect increased percentage of FOXP3+ cells within the iTregs in bexarotene-induced conditions, as shown in Fig. 1B. Overall, these data confirm that iTregs induced in the presence of bexarotene are functionally suppressive.

**Figure 2.**
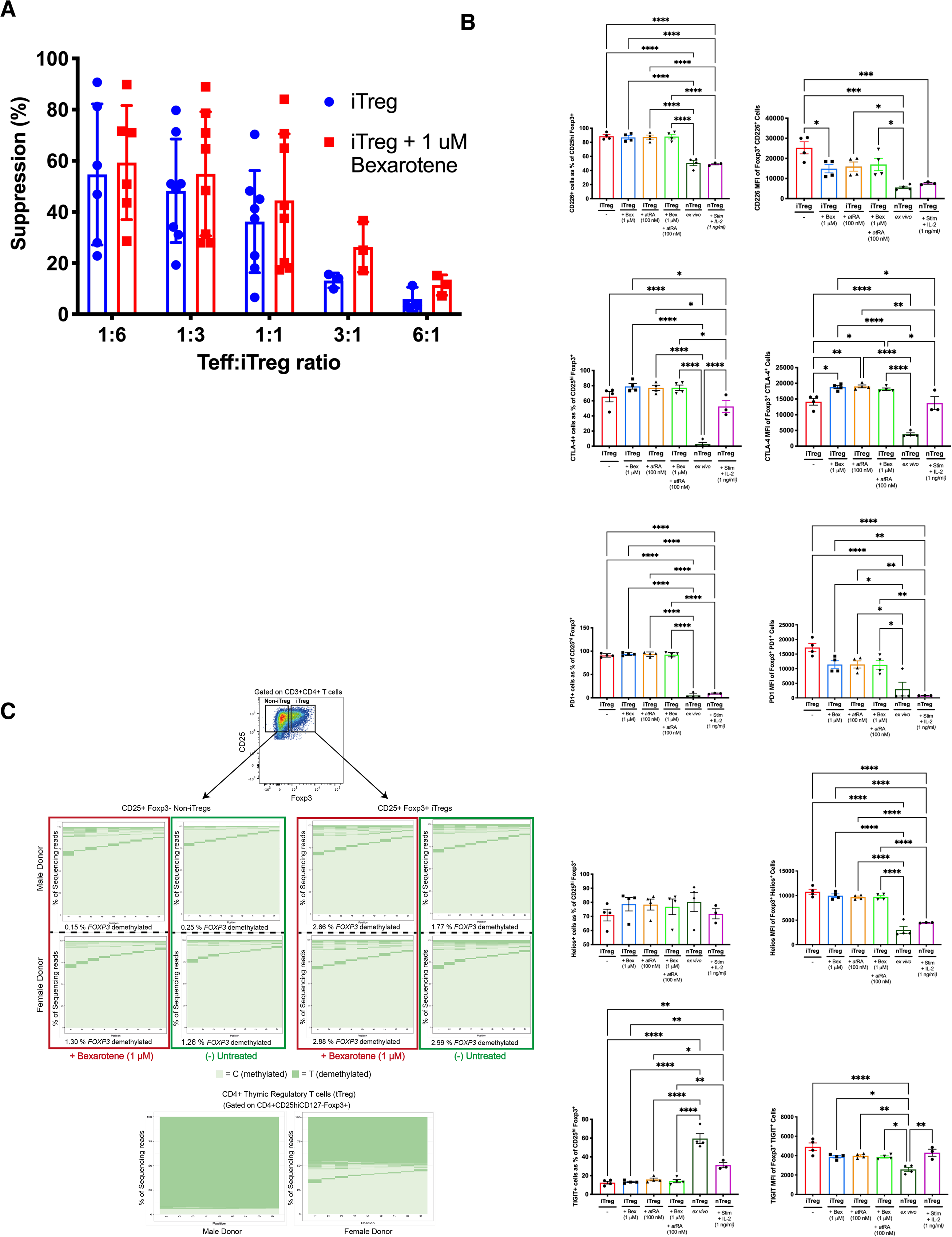
Comparison of iTregs vs donor-matched nTregs. (A) Percent suppression of CD3/CD28 stimulated Teff in the presence of iTregs or iTregs induced in the presence of bexarotene (two-tailed ANOVA, ns). (B) Phenotypic characteristics of iTregs, induced +/−bexarotene, +/−atRA or both vs. donor-matched nTregs stained immediately (ex-vivo) or following culture for 7 days (5 μM CD3, 1 μM CD28, 1 ng/ml IL-2). Shown are mean values ± SEM. iTreg, n=4. ex vivo iTreg, n=4. nTreg + stim, n=3. Significance was calculated with one-way ANOVA, *p<0.05, **p<0.01, **p<0.001, ****p<0.0001 (Tukey’s correction). (C) FOXP3 TSDR methylation, determined by bisulphite sequencing, of iTregs and iTregs induced in the presence of bexarotene vs. donor-matched nTregs. Owing to X inactivation FOXP3 TSDR demethylation is ~50% lower in females compared to males. Plots from two representative donors (one male, one female) are shown; total donors n=6.

### Phenotypic differences between nTregs and iTregs induced in different conditions

To explore any phenotypic differences between human nTregs and iTregs induced under different conditions, the expression of several Treg signature molecules was analysed in iTregs, induced in the presence of *αt*RA or bexarotene versus nTregs (*ex vivo*, unstimulated and stimulated) (Figure 2B).

The proportion of cells expressing the inhibitory receptors CTLA4 and PD1 was higher in iTregs compared to ex vivo nTregs (74.72% vs 2.76% and 92.71% vs 16.56% respectively, Fig 2A). Furthermore, density of CTLA4 and PD1 expression was more than four-fold higher higher in iTregs (MFI: 17,512 vs 3805.5 and 12,857 vs. 3005.0 respectively). Following nTregs stimulation [dr2] surface CLTA4 expression increased significantly [dr3] but remained slightly lower than iTregs induced [dr4] in the presence of bexarotene and/or *αt*RA. In contrast, nTreg PD1 expression remained very low post-stimulation.

Comparing between iTreg conditions, both bexarotene and *αt*RA increased CTLA4 expression (MFI of positive cells) but had no effect on PD 1 expression.

Compared to nTregs (ex vivo and stimulated), fewer iTregs expressed the inhibitory molecule TIGIT (59.48%, 41.11% and 19.42%, respectively), but those expressing TIGIT did so at a higher density (MFI 4914.8 vs 2575.25; p=<0.0001). However this difference was lost when nTregs were stimulated. In contrast, expression of the T-cell activating molecule CD226, which competes against TIGIT for binding to their shared ligands (CD155 and CD122) was found to be increased on iTregs compared to ex vivo and activated nTregs, both in terms of percent expression and MFI. Treg induction in the presence of bexarotene, *αt*RA or both reduced TIGIT (p=0.034, p=0.048, p=0.040) and CD226 (p=0.002, p=0.004, p=0.008) expression on a per-cell basis, but did not affect the percent positive.

A higher percentage of bexarotene or *αt*RA induced iTregs expressed HELIOS, 83.68% and 81.40%, respectively versus 70.975% in untreated iTregs, this increase was not observed in cells cultured with both compounds, or DMSO control (as both bexarotene and *αt*RA are reconstituted in DMSO). bexarotene and *αt*RA had no effect upon the per-cell expression of HELIOS.

### Bexarotene does not alter the *FOXP3TSDR* of iTregs

To address the question of iTreg lineage stability, we assessed methylation status of the FOXP3 Treg specific demethylation region (TSDR) within intron 1 of *FOXP3* using bisulfite sequencing (Rainbow *et al* 2015). Tregs with a highly demethylated TSDR are known to exhibit stable FOXP3 expression whereas cells that transiently express FOXP3 are highly methylated within this region and may lose Treg associated activity. As *FOXP3* is on the X-chromosome, X-chromosome inactivation approximately halves the level of TSDR demethylation seen within female donors.

As expected, flow sorted nTregs were fully demethylated whereas iTregs displayed a methylated *FOXP3* TSDR, comparable to that of FOXP3-cells from the same culture. The addition of 1 μM bexarotene had no effect on the TSDR methylation of iTregs (Figure 2C).

### Bexarotene suppresses the induction of Th17 cells

In addition to enhancing the conversion of FOXP3+ iTregs *in vitro* from naive CD4+ T cells, RXR agonists have been reported to inhibit murine Th17 development *in vitro* and *in vivo* in an RAR independent manner. Here we sought to determine whether bexarotene was capable of suppressing human Th17 induction in a similar manner.

Briefly, MACS separated naive CD4+ lymphocytes stimulated in serum free media, with or without the addition of bexarotene and/or *αt*RA, under Th17 skewing conditions (namely in the presence of TGF-⍰, IL-6, IL-1B, IL-23 and with neutralising antibodies against IL-4 and IFN-⋁).

Both bexarotene and *αt*RA were equally able to suppress human Th17 cell differentiation *in vitro*, as measured by intracellular IL-17A cytokine staining and IL-17A secretion (through ELISA of supernatant) (Figure 3B, C). No synergistic effect was observed, with bexarotene and *αt*RA together suppressing Th17 cell differentiation to the same degree as found with either bexarotene or *αt*RA alone.

**Figure 3.**
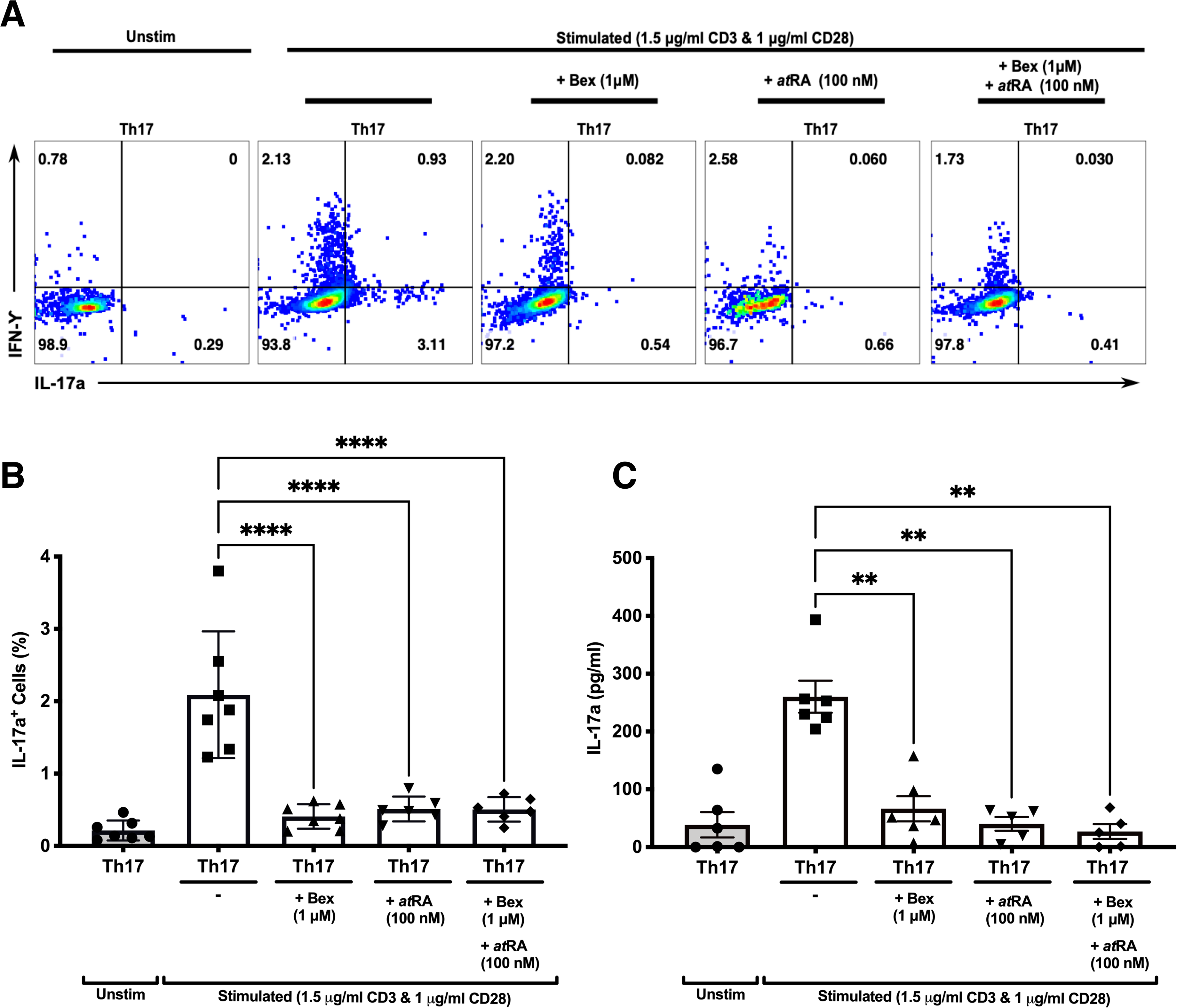
Bexarotene significantly reduces the differentiation of CD4+ naive lymphocytes into Th17 IL-17a+ cells in vitro. (A) Representative dot plots showing intracellular Th17 and IFN-γ following culture of human naïve CD4 cells in Th17 skewing conditions +/− bexarotene, +/−atRA or both. (B) Summary data showing percent IL-17+ cells as a percent of total live CD4 cells at the end of culture (n=7). (C) Supernatant IL-17a as measured at day 7 (n=7). Shown are mean values ± SEM. Significance was calculated with repeated measures one-way ANOVA. Šidák’s correction was performed. *p<0.05, **p<0.01, ***p<0.001, ****p<0.0001.

These findings demonstrate for the first time that an RXR specific agonist can actively inhibit human Th17 differentiation and the secretion of IL-17A following stimulation *in vitro*.

### No differences are observed in the peripheral blood of individuals treated with bexarotene on the CCMR-One study

Patients enrolled within the CCMR One trial receiving bexarotene were immunophenotyped to examine if the immunomodulatory effects demonstrated *in vitro* could be replicated *in vivo*. PBMCs were collected from patients randomised to receive bexarotene pre-and at months 2, 4 and 6 of treatment. Serum samples were collected at the same time points for measurement of IL-17A.

The frequency of CD4+ T cells was unchanged by treatment (Figure 4A), and no difference was found in the frequency of circulating CD4+ Tregs (Figure 4B, C), Th17 cells (Figure 4G,H) (identified by intracellular flow cytometry) or serum IL-17A levels (Figure 4I) in patients treated with bexarotene. Furthermore, in line with our *in vitro* data, no difference was found in the methylation status of Tregs from both male and female donors (Figure 4D, F).

**Figure 4.**
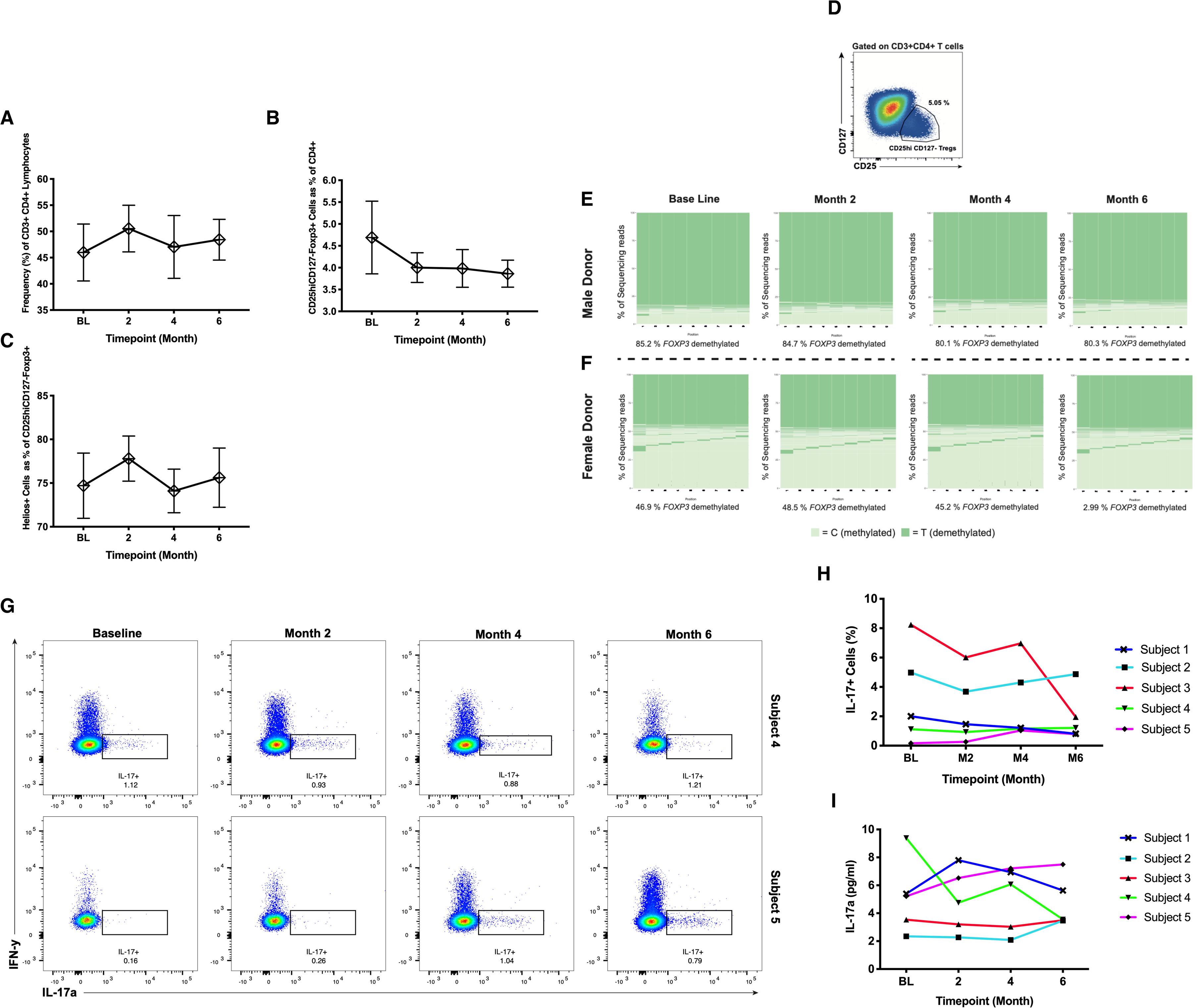
Peripheral blood immune phenotyping of patients randomised to receive bexarotene on the CCMR-One trial. Samples were analysed at baseline, then at months 2, 4 and 6 following treatment. (A) Frequency of CD3+CD4+ cells as a percent of total lymphocytes. (B) Frequency of CD25hiCD127-FOXP3+ Tregs as percent of CD4+ cells. (C) Frequency of Helios+ cells as a percent of CD25hiCD127-FOXP3+ Tregs. Shown are mean values ± SEM, n=5. (D, E, F) FOXP3 TSDR methylation of flow sorted, circulating CD4+CD25hi CD127-Tregs pre and post bexarotene (n=5; two representative donors are shown). (G) intraceullular IL17 and IFN-γ in CD4+ cells, from patients pre and post bexarotene, following stimulation with PMA + Ionomycin + Brefeldin A. Dot plots from two representative donors are shown. (H) Subjects 1 - 5 percent IL-17a+ cells and (I) IL-17a supernatant concentration. Each sample was run in duplicate.

## Discussion

We have demonstrated that the RXR agonist bexarotene, which has recently been shown to promote remyelination in individuals with MS has immunomodulatory effects on human cells-promoting naive CD4^+^ T-cell differentiation into Tregs and suppressing their differentiation into Th17 cells. Our findings lend support to the development of RXR agonists as a therapy for MS.

Despite opposing functions, Th17 cells and iTregs share a closely related developmental pathway mediated by TGF-β. In the presence of TGF-β and IL-6 or IL-21 (produced by activated T-cells and DCs) naive CD4+ cells differentiate into Th17 cells. However, in the absence of proinflammatory cytokines, and in the presence of TGF-β and IL-2, naive cells differentiate into Tregs. TGF-β induces SMAD2 and SMAD3 phosphorylation which in turn activate FOXP3, which drives cells towards the Treg lineage (in contrast IL-6 inhibits SMAD signalling). In addition, IL-2 phosphorylates STAT5 which activates FOXP3. iTregs induced *in vivo* are termed “pTregs” (for peripherally induced), to distinguish them from thymically derived Tregs (also known as naturally occuring or “nTregs”). It is thought that pTregs may constitute a significant proportion of Tregs in the periphery particularly at certain sites such as the lamina propria and maternal placenta where they may maintain tolerance against commensal bacteria and the developing fetus [7]. In addition to their shared dependency on TGF-β, many studies have reported plasticity between iTregs and Th17 cells under certain conditions. For example, studies tracing the fate of IL-17^+^ cells in the gut, reveal that many at some point express FOXP3 [8]. Given this, it is not surprising that the Th17/Treg balance has been shown to play a critically important role in many autoimmune diseases, including MS [9].

In contrast to published murine studies, we have shown that bexarotene promotes human iTreg differentiation independent of RAR signalling. Therefore RXR supplementation alone should be sufficient to promote iTreg induction in patients. Independence from RAR signalling was shown both by culturing cells in serum-free, and therefore *αt*RA free media, and by using the pan-RAR antagonist AGN193109. Also counter to murine data is our observation that bexarotene and *αt*RA do not work synergistically to enhance human iTreg differentiation in vitro. One possibility is that *αt*RA was used at concentrations reaching the maximum threshold for atRA enhancement of human iTregs differentiation. *αt*RA is thought to promote iTreg differentiation through enhancing TGF-β driven SMAD3 signalling and inhibiting the activity of IL-6 and IL-23 [10], meaning it is possible that these mechanisms were saturated leaving bexarotene unable to promote further *αt*RA-mediated enhancement. In addition to RXR forming a heterodimer with RAR, it can also form a homodimer or dimerise with alternative binding partners - including the 1a,25-(OH)2 vitamin D3 receptor (VDR), peroxisome proliferator-activated receptor (PPAR), liver X receptors (LXR), thyroid hormone receptors (TR), pregnane X receptors (PXR) and farnesoid X receptor (FXR) [11].

We have not determined through which receptor-dimer complex bexarotene promotes iTreg differentiation. However, as it occurs independently of RAR engagement it must do so either through RXR/RXR homodimers, or via permissive nuclear receptors, such as PPAR or LXR. Evidence in support of PPAR engagement comes from studies showing that PPAR-α and PPAR-γ agonists also promote FOXP3 expression and TSDR demethylation in CD4+CD25-T cells upon TCR stimulation in the presence of TGF-β and IL-2 [12].

Compared to thymically derived nTregs (ex vivo or activated), iTregs (induced in all conditions, including bexarotene) expressed higher surface levels of the inhibitory molecules CTLA-4 and PD-1 (both in terms of percentage and on a per-cell basis). Per-cell surface CTLA-4 expression was further increased by exposure to bexarotene or *αt*RA. Cytotoxic T-lymphocyte-associated protein 4 (CTLA-4) is a crucial inhibitory molecule and homolog to the co-stimulatory molecule CD28, with which it shares ligands (CD80 and CD86 expressed on APCs). Binding of CD80/86 (on antigen presenting cells) to CD28 on T-cells provides a costimulatory signal, whereas CTLA-4 captures CD80/68 from the surface of APCs by transendocytosis, preventing CD28 engagement and T-cell activation [16]. PD-1 interactions with PD-L1 and PD-L2 also down-modulate T-cell immune responses [17]. In contrast to CTLA-4 and PD-1, significantly fewer iTregs expressed the co-inhibitory molecule TIGIT (T cell Ig and ITIM domain) compared to nTregs, although iTregs expressing TIGIT did so at a higher level. Expression of the co-stimulatory molecules CD226 was also increased on iTregs (percent and per-cell expression) compared to nTregs, although per-cell expression was slightly lower when Tregs were induced in the presence of bexarotene and/or *αt*RA. Similar to CD28 and CTLA-4, CD226 and TIGIT share and compete for ligands expressed on APCs (CD112 and CD155) promoting or inhibiting T-cell activation respectively [18, 19]. Despite high CD226:TIGIT expression, and perhaps as a consequence of high PD-1 and CTLA-4 expression, iTregs induced in the presence of bexarotene were found to be at least as suppressive *in vitro* as iTregs induced in standard conditions.

In addition to promoting iTreg differentiation, we have shown that bexarotene is also capable of suppressing human Th17 development *in vitro*, as measured through intracellular IL-17A staining and changes in supernatant IL-17A. This is the first report using human cells and is keeping with murine studies where RXR ligands have been shown to suppress Th17 development. Two such studies report that RXR ligands are able to suppress Th17 development, attenuate active and Th17 mediated passive EAE and suppress the numbers of CD4+ lymphocytes capable of secreting pro-inflammatory cytokines [3, 5]. Furthermore, RXR ligands have been shown to reduce the expression of cell activation markers Ki-67 and CCR6 on Th17 cells which is required for entry into the central nervous system (CNS). These results are encouraging for diseases such as MS, thought to be driven by aberrant regulation of Th17 cell responses that are found with increased frequency in the periphery and the CNS of patients [13, 14].

In this study, we took advantage of access to blood samples from individuals with MS receiving bexarotene as participants on the CCMR-One phase 2a trial, to explore the effects of bexarotene *in vivo*. Disappointingly our *in vitr*o observations were not mirrored in the blood of individuals receiving bexarotene. This is perhaps unsurprising given that we would expect bexarotene to exert its effects at sites of CD4 naive cell activation and differentiation, which would be unlikely to affect circulating Treg/Th17 numbers. Our results are also in keeping with findings with use of bexarotene in cutaneous T cell lymphoma (CTCL), where no change in the number of circulating Tregs was found in peripheral blood [15], although in this study “Tregs” were defined purely on the basis of CD25hi expression.

Bexarotene was poorly tolerated in CCMR-One. Of the 32 patients dosed in Cambridge, 16 developed central hypothyroidism and 14 developed elevated triglycerides, both well recognised side-effects of bexarotene. Given this, it is unlikely that bexarotene will progress further into clinical development as a therapeutic strategy in MS. However, these results do provide hope and momentum behind the development of other RXR-isoform specific agonists.

In summary we suggest that RXR agonism may be an effective therapeutic approach for the treatment autoimmune disorders particularly those such as MS where there is an imbalance in the Treg/Th17 axis. MS is a particularly attractive target as RXR agonists have also been reported to promote remyelination.

## Methods and Materials

### Ethics Statement

Peripheral blood mononuclear cells (PBMCs) were isolated from anonymous healthy donor buffy coats, which were obtained from the Addenbrooke’s Hospital NHS blood and transplant service (Cambridge, United Kingdom). All individuals gave written consent, and the study was approved by a local ethical review committee (REC: 11/EE/0007). Individuals with relapsing remitting multiple sclerosis treated on the CCMR-One study gave blood for research under REC 15/LO/0108.

### Cell preparation

Human PBMCs were isolated from the fresh buffy coats of whole blood of healthy donors by Ficoll-Paque Plus centrifugation (GE Healthcare, Sweden). PBMCs were counted by trypan blue exclusion and resuspended in PBS. Naive CD4+ T cells were either isolated through magnetic activated cell sorting (MACS) or fluorescence-activated cell sorting (FACS). With MACS, naive T cells were isolated through negative selection from total PBMC using the naive CD4+ T Cell Isolation Kit, human (Miltenyi, 130-094-131), according to the manufacturer’s instructions. With FACS, naive CD3+CD4+CD127+CD25lowCD45RA+CD62L+CD27+CCR7+ were isolated from CD4+ enriched PBMCs following negative isolation using CD4+ T cell Isolation Kit, human (Miltenyi, 130-096-533), using a BD Influx (supplemental figure S1). Staining for FACS was done on cell surface markers, live/dead, CD3, CD4, CD127, CD25, CD62L, CD45RA, CD27 and CCR7 with relevant mAbs (BioLegend and BD). CD25^high^ “nTregs" and CD3+CD4+CD127+CD45RA-CD62L-“Teffs” were also isolated from the same CD4+ enriched population and were cryopreserved in 10% DMSO (Sigma-Aldrich) and 90% FBS (Life Technologies, Thermo Fisher Scientific), for later use in suppression assays. Naive CD4+ T cells were cultured at 5% CO_2_/37°C in serum-free X-Vivo 15 medium (Lonza).

### iTreg differentiation

Naive CD4+ T cells cells were plated under iTreg differentiation conditions at 1.0×10^5^ cells/well in 96 U-bottom well plates (Falcon). For cell stimulation, plates were coated at least 5 hours prior to use with 5 ug/ml plate-bound anti-CD3 antibody (clone OKT3; BioLegend, LEAF grade), which is then washed off with PBS, with 1 ug/ml soluble anti-CD28 antibody (BioLegend, LEAF grade) and 5 ng/ml IL-2 (carrier-free; Tocris R&D Systems) added with cells. Cells treated with only these functioned as the mock control. For iTreg induction, TGF-*β* (5ng/ml; carrier-free; Tocris R&D Systems), anti-IFN*γ* antibody (2 ug/ml; clone MD-1, Biolegend), bexarotene (1 *μ*M, Abcam) or *αt*RA (100 nM; Sigma-Aldrich) were added additionally. The DMSO control (for bexarotene, *αt*RA and AGN190) had no effect on FOXP3 expression. Cells were incubated for 7 days unless otherwise stated.

### Th17 cell differentiation

After either MACS or FACS isolation, naive cells were plated under Th17 differentiation conditions at 5×10^5^ cells/well in 96 U-bottom well plates (Falcon). For cell stimulation, plates were coated at least 5 hours prior to use with 3 ug/ml plate-bound anti-CD3 antibody (clone OKT3; BioLegend, LEAF grade), 1 ug/ml soluble anti-CD28 antibody (BioLegend, LEAF grade) and 2 ng/ml IL-2 (carrier-free; Tocris R&D Systems). Cells treated with only these functioned as the mock control. For Th17 induction, TGF-*β* (30 ng/ml; carrier-free; Tocris R&D Systems), IL-6 (30 ng/ml; carrier-free; Tocris R&D Systems), IL-1*β* (10 ng/ml; carrier-free; Tocris R&D Systems), IL-23 (50 ng/ml; carrier-free; Tocris R&D Systems), anti-IFN*γ* antibody (2 ug/ml; clone MD-1, Biolegend), anti-IL-4 antibody (2 ug/ml; clone MD-1, Biolegend), bexarotene (1 *μ*M, Abcam), or *αt*RA (100 nM; Sigma-Aldrich) were added additionally. The DMSO control (for bexarotene, *αt*RA and AGN190) had no effect on IL-17A cytokine expression. Cells were incubated for 7 days unless otherwise stated.

### In vitro suppression assays

For use in suppression assays, on day 7 iTregs were washed twice in PBS and their viability and count assessed with trypan blue exclusion. Cells were then allowed to rest for 3 hours in fresh, serum free X-Vivo 15 media prior to addition into suppression assay cell culture at 5% CO_2_ at 37°C. The mock IL-2 treated control cells were also washed, counted and rested in the same manner as for iTregs. Cryopreserved Teffs from donors were thawed quickly within a water bath at 37°C, washed and rested for 4 hours at 5% CO_2_ 37°C. To discriminate between Teffs and Tregs in the suppression assay, Teffs were labelled with 5 mM cell proliferation dye EFV450, and iTregs were labelled with 5 mM cell proliferation dye BV670 (both from eBioscience). Teffs were plated at 5×10^4^ cells per well in duplicate, with and without iTregs, in serum free X-Vivo 15 media alone. Cells were cultured for 3 days. For the final analysis, dead cells were excluded using dead cell exclusion staining (Zombie NIR; eBioscience), and cells were counterstained with Abs to CD4. Due to unknown suppressive capacity, iTregs were titrated up so that the iTreg/Teff ratio was 1:1 to 3:1 and 6:1. Duplicate control wells of CD4+ T cells without stimulus, and Tregs, at various dilutions, with and without stimulus, were also cultured. Treg Suppression Inspector beads (Miltenyi Biotec) were used to stimulate the assay, according to the manufacturer’s instructions.

### Flow cytometry and antibodies

#### Viability staining

Cells taken from culture were washed once with 1 mL PBS and resuspended in 100 ul PBS. Cells were then stained with LIVE/DEAD™ Fixable Aqua Dead Cell Stain Kit (Thermo Fisher Scientific) for 30 minutes, in the dark at 4°C. Cells were then washed twice with 1 mL of FACS buffer (0.5% BSA/PBS) and taken for subsequent cell surface marker staining.

#### Surface staining

Cell surface staining was performed in the dark at 4°C in antibody dilutions in FACS buffer for 30 minutes, with a final staining volume of 100 ul. Cells were then washed twice with 1 mL PBS and either taken for flow cytometry analysis or used for subsequent intracellular staining

### Cell permeabilisation and intracellular Staining

Cell permeabilisation and fixation was performed using the Foxp3 Staining Buffer Set (eBioscience), according to the manufacturer’s instructions at room temperature for 40 minutes. Intracellular staining was then performed, according to manufacturer’s instructions in the dark at 4°C. To measure intracellular cytokines, cells were stimulated for 4h prior to staining with 1x Cell Activation Cocktail (PMA, ionomycin, and Brefeldin A; 500X).

### Antibodies (anti-human)

The complete list of antibodies used is as follows and were purchased from eBioscience, BD Biosciences or BioLegend, unless otherwise indicated. Brackets indicate (fluorochrome, antibody clone).

#### Sorting panel

CD45RA (AF488; HI100), CCR7 (PerCP-Cy5.5; G043H7), Zombie Aqua (live/dead stain), CD27 (BV605; 323), CD3 (BV650; UCHT1), CD62L (BV786; DREG-56), FOXP3 (PE; 259D, 259D/C7, 206D), CD127 (PE-Cy7; eBioRDR5), CD25 (APC; M-A251) and CD4 (APC-R700; RPA-T4).

#### iTreg staining

FOXP3 (PE; 259D), CD25 (APC, 2A3), CD3 (BV650; UCHT1), CD4 (APC-R700, RPA-T4), CD127 (PE-Cy7; eBioRDR5).

#### Th17 staining

CD3 (BV650, UCHT1), CD4 (APC-R700, RPA-T4), CD25 (APC, 2A3), CD127 (PE-Cy7, eBioRDR5), IFN-y (BUV395; B27), IL-17 (PE, SCPL1362) IL-21 (BV421; 3A3-N2.1).

#### CCMR-One

FOXP3 (PE; 259D), CD25 (BV421, 2A3 and MA251), CCR6 (BV650, 11A9), CCR4 (BB700, 1G1), CD3 (BV605, SK7), CD45RA (BV785, HI100), HELIOS (FITC, 22F6), CD8 (PE-Cy5, RPA-T8), CD4 (APC, RPA-T4), HLA-DR (AFR-700, L243), T.S. monocyte blocker, CD127 (PE-Cy7, PCH101), CD27 (APC-eFluor 780, 0323), CCR4 (PerCP eFluor 710, D8SEE).

### Acquisition

Acquisition was performed on a BD LSR Fortessa and compensation was performed manually using single stained colour controls. Cell sorting was performed on BD Influx under sterile conditions. FACS data was analysed using FlowJo (V10.1) and sort reports were collected for each cell sort experiment.

### Enzyme-linked immunosorbent assay (ELISA) measurements

IL-17a ELISAs were performed using Human IL-17A (homodimer) ELISA Ready-SET-Go!™ Kit (Invitrogen™; eBioscience™), according to the manufacturer’s instructions. Plates were coated the day prior to beginning the ELISA and kept covered at 4°C.

### Analysis of TSDR methylation

CD4+ CD25+ Foxp3+ iTregs and CD4+ CD25+ Foxp3-non-Tregs were sorted by FACS from “bulk iTreg cultures” from 4 male and 4 female donors, then stored immediately as a dry frozen pellet for bisulphite sequencing. DNA of FACS sorted iTregs and control cells from female and male donors was isolated using the DNeasy Blood and Tissue Kit (Qiagen), according to the manufacturer’s instructions. Bisulphite sequencing was performed in house, as previously described (Rainbow *et al* 2015). Six replicates of 3000 cells per replicate (5 ng DNA) were analysed from a single donor and the median (with range) reads with eight or nine sites demethylated at FOXP3 for the replicates, is reported.

### Statistical analysis

All flow cytometry data were analysed in-house with FlowJo (v10.1); all other data analyses and statistical fitting were performed in-house with GraphPad (Prism). Where appropriate, ANOVA and two-tailed t-tests were performed to assess for differences in Foxp3 protein expression between wells and experiments. One-way ANOVAs were corrected using Tukey’s multiple comparisons test. Repeated measures one-way ANOVAs and two-tailed t-tests were corrected using Šidák’s correction. The test and corresponding correction performed is indicated in each figure legend.

## Acknowledgements

This research was supported by the Wellcome Trust (RG79413), Cambridge NIHR BRC Cell Phenotyping Hub and NIHR Cambridge Biomedical Research Centre (BRC-1215-20014). The views expressed are those of the author(s) and not necessarily those of the NIHR or the Department of Health and Social Care. CG was funded by a BBSRC CASE studentship. LJ and JJ were funded by the Wellcome Trust (RG79413).

## Notes

### Competing Interest Statement

The authors have declared no competing interest.

