## Supplemental 1 for "The MS remyelinating drug bexarotene (an RXR agonist) promotes induction of human Tregs and suppresses Th17 differentiation *in vitro*"

### Supplemental data

**S1.** T cell sort gating protocol

#### T Cell Sort Gating Protocol

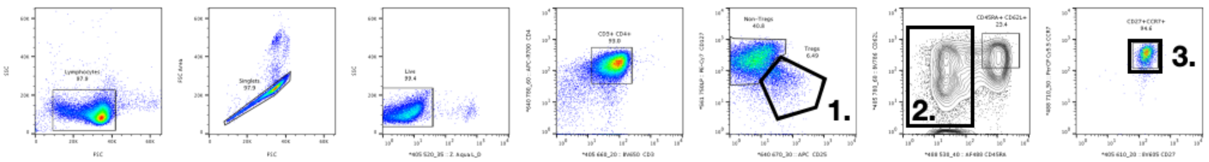


#### Sorting Panel

| 530/30 | **CD45RA** | **AF488** |
|---|---|---|
| 695/40 | CCR7 | PerCP-Cy5.5 |
| 525/50 | Zombie Aqua | BV510 |
| 605/12 | CD27 | BV605 |
| 650/8 | CD3 | BV650 |
| 780/60 | CD62L | BV786 |
| 585/15 | FOXP3 | PE |
| 780/60 | CD127 | PE-Cy7 |
| 670/14 | CD25 | APC |
| 730/45 | CD4 | APC-R700 |

**1. nTregs**
**(CD3+ CD4+ CD25hi CD127-)**

**2. Teffs**
**(CD3+ CD4+ CD127+ CD45RA-)**

**3. CD4+ Naive T cells**
**(CD3+ CD4+ CD127+ CD45RA+ CD62L+ CCR7+ CD27+)**
